# A data-driven investigation of human action representations

**DOI:** 10.1101/2022.09.22.509054

**Authors:** Diana C. Dima, Martin N. Hebart, Leyla Isik

**Affiliations:** Dept. of Cognitive Science, Johns Hopkins University, Baltimore, USA; Dept. of Computer Science, Western University, London, Canada; Vision and Computational Cognition Group, Max Planck Institute for Human Cognitive and Brain Sciences, Leipzig, Germany

## Abstract

Understanding actions performed by others requires us to integrate different types of information about people, scenes, objects, and their interactions. What organizing dimensions does the mind use to make sense of this complex action space? To address this question, we collected intuitive similarity judgments across two large-scale sets of naturalistic videos depicting everyday actions. We used cross-validated sparse non-negative matrix factorization (NMF) to identify the structure underlying action similarity judgments. A low-dimensional representation, consisting of nine to ten dimensions, was sufficient to accurately reconstruct human similarity judgments. The dimensions were robust to stimulus set perturbations and reproducible in a separate odd-one-out experiment. Human labels mapped these dimensions onto semantic axes relating to food, work, and home life; social axes relating to people and emotions; and one visual axis related to scene setting. While highly interpretable, these dimensions did not share a clear one-to-one correspondence with prior hypotheses of action-relevant dimensions. Together, our results reveal a low-dimensional set of robust and interpretable dimensions that organize intuitive action similarity judgments and highlight the importance of data-driven investigations of behavioral representations.

## Introduction

Our ability to rapidly recognize and respond to others’ actions is remarkable, given the wide variety of human behaviors that span different contexts, goals, and motor sequences. When we see a person performing an action, we integrate visual information, social cues and prior knowledge to interpret the event, a task that still challenges state-of-the-art machine learning algorithms. How does the mind make sense of this complex action space?

Previous work on action understanding in the mind and brain has focused on hypothesis-driven efforts to identify critical action features and their neural underpinnings. This work has highlighted semantic content (Lingnau & Downing, 2015; Tucciarelli et al., 2019), social and affective features (Dima et al., 2022; Tarhan & Konkle, 2020; Wurm et al., 2017), and visual features (Tarhan & Konkle, 2020; Wurm & Caramazza, 2019) as essential components in visual action understanding. However, such an approach requires the experimenter to pre-define actions and their potential organizing dimensions, necessarily limiting the hypothesis space. Action categories have commonly been defined based on the verbs they represent (Bedny & Caramazza, 2011) or everyday action categories as listed, for example, in the American Time Use Survey (ATUS; Dima et al., 2022; Tarhan et al., 2021; Tarhan & Konkle, 2020). Given the diversity of actions, a low-dimensional, flexible representation may be a more efficient way to organize them in the mind and brain; but generating the hypotheses that could uncover this representation remains difficult, especially for naturalistic stimuli that vary along multiple axes.

Data-driven methods provide an alternative to pre-defined representational spaces and have achieved great success in mapping perceptual and psychological representations in other visual domains. In object recognition, a data-driven computational model revealed 49 interpretable dimensions capable of accurately predicting human similarity judgments (Hebart et al., 2020). Recent work has extended this method to near scenes, known as reachspaces, and identified 30 dimensions capturing their most important characteristics (Josephs et al., 2021). Low-dimensional representations have been also proposed that explain how people perceive others and their mental states (Gray et al., 2007; Thornton & Tamir, 2020) or psychologically meaningful situations (Parrigon et al., 2016; Rauthmann et al., 2014).

To date there has been only limited data-driven work in the action domain. Using principal component analysis (PCA) of large-scale text data, a low-dimensional taxonomy of actions has been shown to explain neural data and human action judgments (Thornton & Tamir, 2021a), as well as guide predictions about actions (Thornton & Tamir, 2021b). However, since this taxonomy was generated from text data, most of these dimensions were relatively abstract (e.g. *creation, tradition, spiritualism*), and it is unclear whether a similar set of dimensions would emerge from visual action representations. In the visual domain, six broad semantic clusters were shown to explain semantic similarity judgments of controlled action images (Tucciarelli et al., 2019), suggesting that actions may be semantically categorized at the superordinate level. However, it remains unclear how this finding would generalize to more natural and diverse stimulus sets.

We analyzed a dataset containing unconstrained behavioral similarity judgments of two sets of natural action videos from the Moments in Time dataset (Monfort et al., 2019) collected in our prior study (Dima et al., 2022). Behavioral similarity has often been used as a proxy for mental representations (Edelman, 1998; Murphy, 2002; Shepard, 1987) and has been shown to correlate with neural representations (Bankson et al., 2018; Charest et al., 2014; Cichy et al., 2019; Proklova et al., 2019; Wardle et al., 2016). Specifically, the perceived similarity of actions has been found to map onto critical action features, such as their goals or their social-affective content, as well as onto the structure of neural patterns elicited by actions (Dima et al., 2022; Tarhan et al., 2021; Tucciarelli et al., 2019).

Here, we employ a data-driven approach, sparse non-negative matrix factorization (NMF; Hoyer, 2004) to recover the dimensions underlying behavioral similarity. We show that a cross-validated approach to dimensionality reduction produces a low-dimensional representation that is interpretable by humans and generalizes across stimulus categories. Importantly, the dimensions recovered by NMF are more robust than those generated by the more commonly used PCA. The non-negativity constraint is known to yield a parts-based description, supporting dimension interpretability (Lee & Seung, 1999).

Using human labeling and semantic embeddings, we find that dimensions map to interpretable visual, semantic, and social axes and generalize across two experiments with different experimental structure, stimuli, and participants. Together, our results highlight the semantic structure underlying intuitive action similarity and show that cross-validated NMF is a useful tool for recovering interpretable, low-dimensional cognitive representations.

## Results

### NMF recovers robust dimensions

We analyzed two datasets consisting of three-second naturalistic videos of everyday actions from the Moments in Time dataset (Monfort et al., 2019). In two previously conducted experiments (Dima et al., 2022), participants arranged two sets of 152 and 65 videos from 18 everyday action categories (ATUS, 2019) according to their unconstrained similarity. The first dataset also included videos of natural scenes as a control category (see Stimuli).

We used sparse non-negative matrix factorization (Hoyer, 2002, 2004) with a nested crossvalidation approach (see Methods) to recover the optimal number of underlying dimensions in the behavioral data (Figure 1). This approach combines sparsity and non-negativity constraints to generate feature embeddings that can capture both categorical and continuous information (see Methods; Hebart et al., 2020; Navarro & Lee, 2004; Zheng et al., 2019). Using only behavioral similarity matrices as its starting point, this method can thus recover interpretable features that may shed light on how actions are organized in the mind.

**Figure 1.**
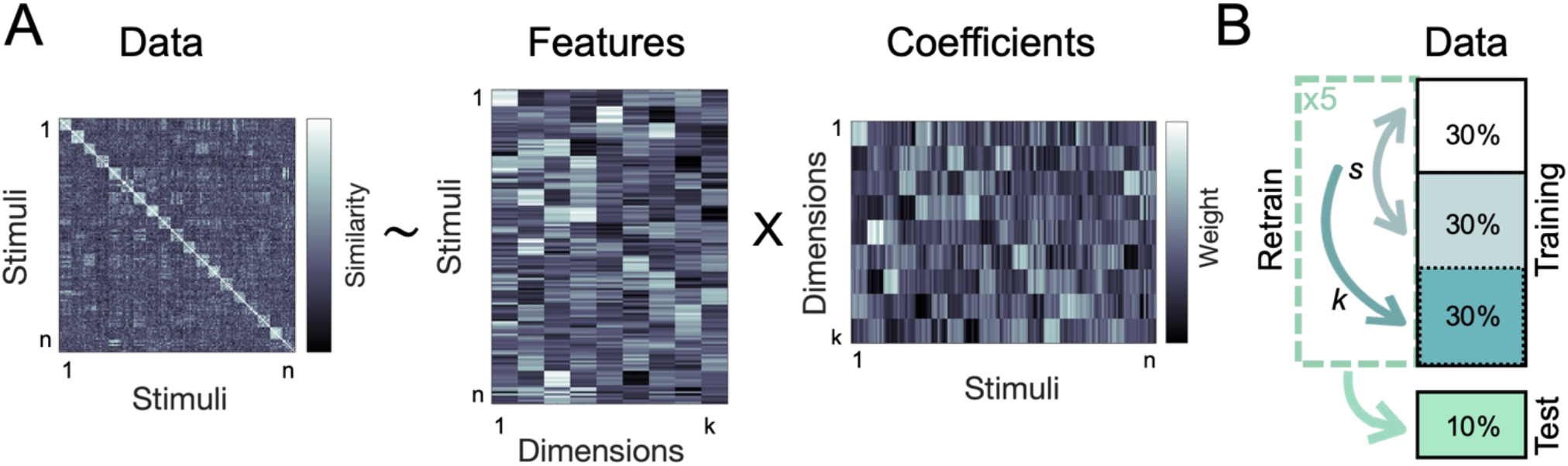
Analysis overview. **A.** Using non-negative matrix factorization, we identified the optimal lower-dimensional approximation of a behavioral similarity matrix. This uncovered the interpretable dimensions underlying the perceived similarity of naturalistic action videos. **B.** NMF cross-validation procedure. Individual similarity ratings were assigned to a cross-validation fold before averaging the input matrices for each fold. The sparsity parameters (*s*) were optimized using two-fold cross-validation on ~60% of the data, with a separate ~30% used to determine the number of dimensions (*k*), and a hold-out set of ~10% used for final evaluation.

Despite differences in stimulus set size and sampling, both experiments were characterized by similar numbers of dimensions (9 and 10 respectively; Supplementary Figure 1) with a sparsity of 0.1. In Experiment 1, the final NMF reconstruction of the entire training set correlated well with the training data (Kendall’s *τ_A_* = 0.46) and the held-out data (*τ_A_* = 0.19, true *τ_A_* = 0.14). Performance was better in Experiment 2, with a training *τ_A_*= 0.75 and a hold-out *τ_A_* = 0.46 (true *τ_A_* = 0.45).

Importantly, the dimensions were robust to systematic perturbations in the underlying stimulus sets (Figure 2). Even after removing critical stimulus categories (such as all outdoor or indoor videos or certain action categories), the NMF procedure resulted in similar numbers of dimensions in both experiments (mean±SD 8.4±0.89 and 8.2±1.64). All dimensions were significantly correlated to those resulting from the full stimulus set, suggesting that the NMF results generalize even after modifying the compositon of the underlying datasets.

**Figure 2.**
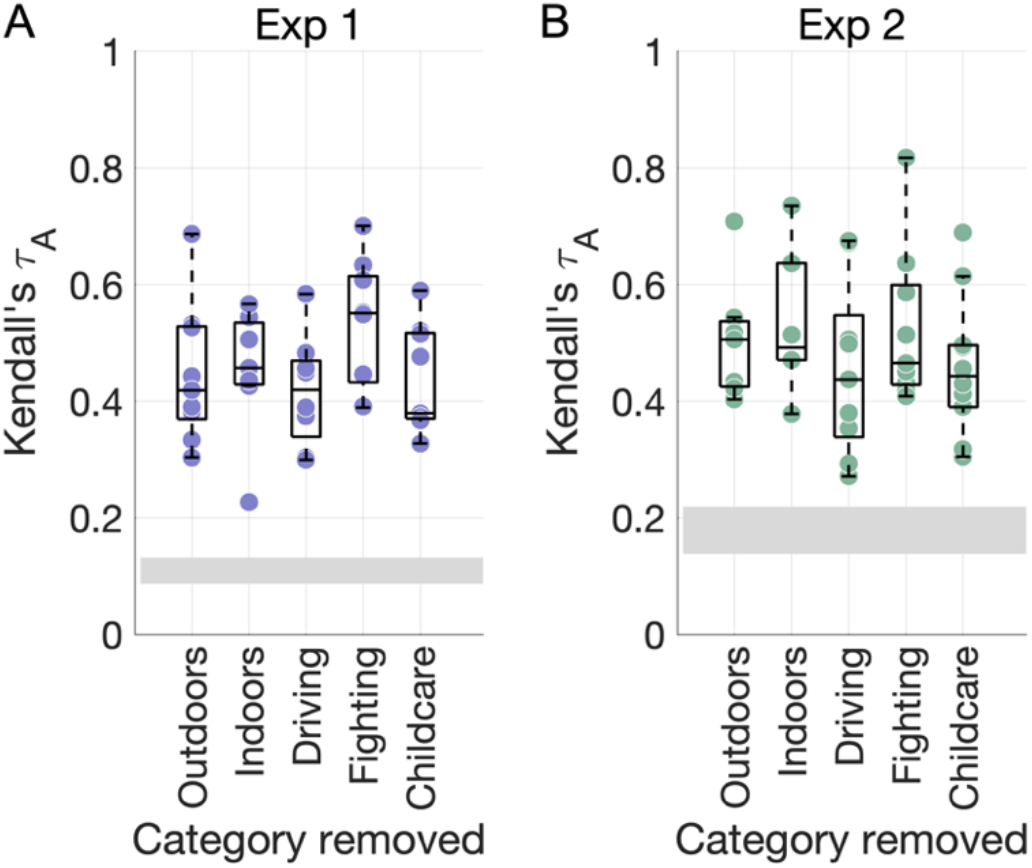
NMF dimension robustness. **A.** The NMF procedure was repeated after removing key stimulus categories from the behavioral RDM from Experiment 1. Each dot shows the maximal correlation between each dimension obtained in the control analysis and any of the original dimensions with the same stimuli removed (repeats allowed). The grey rectangle depicts the chance level (min-max range). **B.** As for **A.**, for Experiment 2.

NMF dimensionality varied less as a function of stimulus set size (average *k* range 6-8.3) than as a function of number of action categories (average *k* range 3.6-10.2; Supplementary Figure 3). Further, NMF dimensions did not map directly onto any single visual, social, or action feature identified in our previous work (Dima et al., 2022; Supplementary Figure 2), suggesting that this method is able to capture additional information not revealed by a hypothesis-driven approach.

Finally, NMF performance was better than that achieved by an equivalent cross-validated analysis using PCA, which recovered 8 dimensions in both experiments (Experiment 1: training *τ_A_* = 0.41, hold-out *τ_A_* = 0.16; Experiment 2: training *τ_A_* = 0.63, hold-out *τ_A_* = 0.41). In the robustness analysis, the number of dimensions generated by PCA after removing critical stimulus categories was less reliable than those obtained with NMF in Experiment 1 (Experiment 1: 7.8±2.49 vs. 8.4±0.98; Experiment 2: 6±1.58 vs 8.2±1.64). While on average correlations with the original dimensions were high, their variance was also more than twice as high as that obtained with NMF (Supplementary Figures 4–5). This suggests that dimensions recovered with PCA are more sensitive to variations in the underlying stimulus set than those found with NMF.

### NMF recovers interpretable dimensions

The hypothesis-neutral dimensions generated by NMF suggest a potential structure to the behavioral space of action understanding. However, further validation is needed to show whether (1) these dimensions are reproducible and (2) to what degree they are interpretable.

To test reproducibility, participants in an online experiment selected the odd video out of a group consisting of seven highly weighted videos and one low-weighted video along each dimension. In a separate online experiment to test interpretability, participants were asked to provide up to three labels for each dimension after viewing the eight highest and eight lowest weighted videos. Their labels were quantitatively evaluated using FastText (Bojanowski et al., 2016), a 300-dimensional word embedding pretrained on 1 million English words.

All dimensions were reproducible in the odd-one-out experiments (Figure 3A; all *P*<0.004), though participants performed significantly better on average in Experiment 1 (mean accuracy 0.8±0.13) than in Experiment 2 (mean accuracy 0.61±0.13, t(15.82) = 3.69, *P*=0.002).

**Figure 3.**
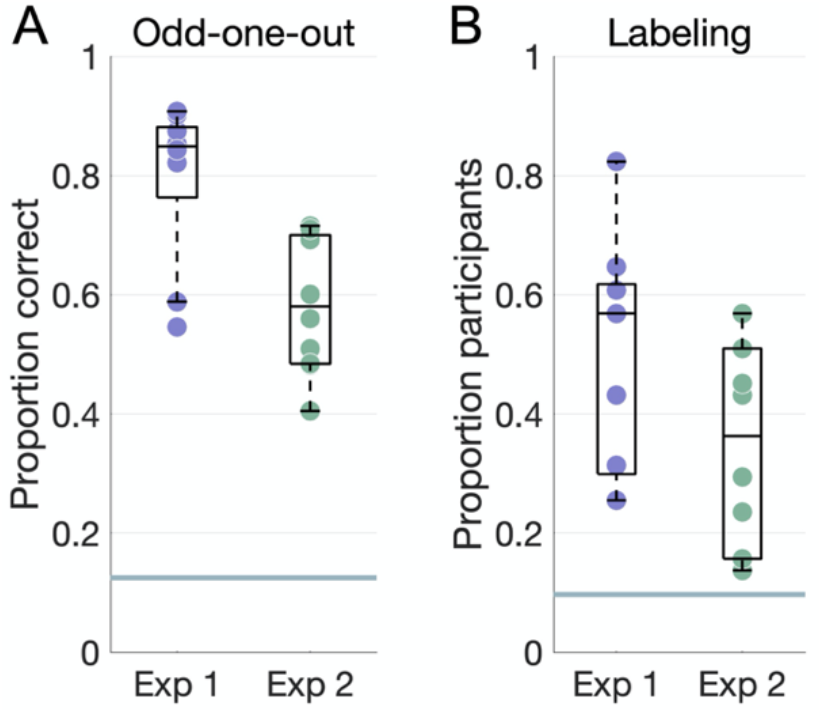
Behavioral results. **A.** Accuracy on the odd-one-out task for each dimension plotted against the chance level of 12.5% (horizontal line). **B.** Proportion of participants who agreed on the top label for each dimension, where agreement is defined as a word embedding dissimilarity in the 10^th^ percentile within all dimensions in both experiments. The horizontal line marks a chance level based on embedding dissimilarity across different dimensions.

Participants’ labels were consistent for most dimensions (Figure 3B). Agreement, as measured via word embeddings, was higher in Experiment 1 (mean proportion 0.5±0.2) than in Experiment 2 (mean proportion 0.34±0.17), though this difference was not significant (t(15.78) = 1.84, *P*=0.08).

The most common labels (Figure 4) captured different types of information, ranging from visual (*nature/outdoors*), to action-related (*eating, cleaning, working*), as well as social and affective (*children/people, talking, celebration/happiness, chaos*). Dimensions in Experiment 2 included more social information overall, with four dimensions labeled with social or affective terms (*talking, people, celebration, chaos*), compared to one in Experiment 1 (*children*). Although many dimensions reflected action categories included in the dataset (*eating, cleaning, working, driving, reading*) or labeled features that explained the most variance our previous experiment (relating to people and affect), the information they provided was richer than the a priori category labels and crossed predefined category boundaries. For example, some videos were highly rated along several different dimensions (e.g. *work* and *learning*), thus capturing the complexity of naturalistic stimuli which often depict several actions or lend themselves to different interpretations.

**Figure 4.**
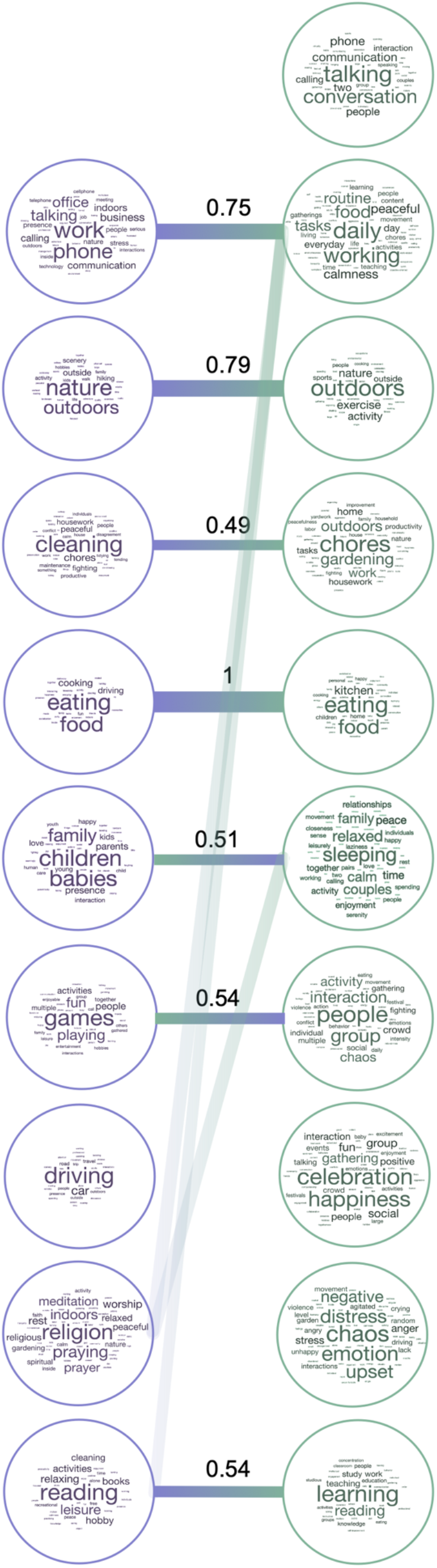
Label correspondence across experiments. Wordclouds showing the labels assigned by participants to each NMF dimension in Experiment 1 (left) and Experiment 2 (right), with larger font sizes representing more frequent labels. Bars connect dimensions from Experiment 1 to their most related dimensions from Experiment 2.

The values shown are normalized relative similarities. Dimensions from Experiment 1 are sorted in descending order of their summed weights, while those from Experiment 2 are organized for clarity of visualization.

Further, not all action categories were reflected in NMF dimensions, suggesting that certain action categories are more important than others in organizing behavior. Certain action categories were absorbed by others (e.g. *eating* included both *eating* and *preparing food*), while other related actions remain separated (e.g. *work* was split into *office work* vs *chores/cleaning*).

### A shared semantic space

To better understand the relationship between dimensions revealed by the two datasets, we calculated Euclidean distance between averaged word embeddings for dimensions in each experiment (see Methods). This analysis revealed several dimensions that were present in both datasets: *eating, nature/outdoors, learning/reading, chores/cleaning*, and *work* (Figure 5). Furthermore, some dimensions were moderately related to several others: *games: people, celebration; work: talking, working; reading: working, learning*. In Experiment 1, the only dimension that did not have a counterpart in Experiment 2 was *driving*, possibly because of the low number of driving videos in Experiment 2.

**Figure 5.**
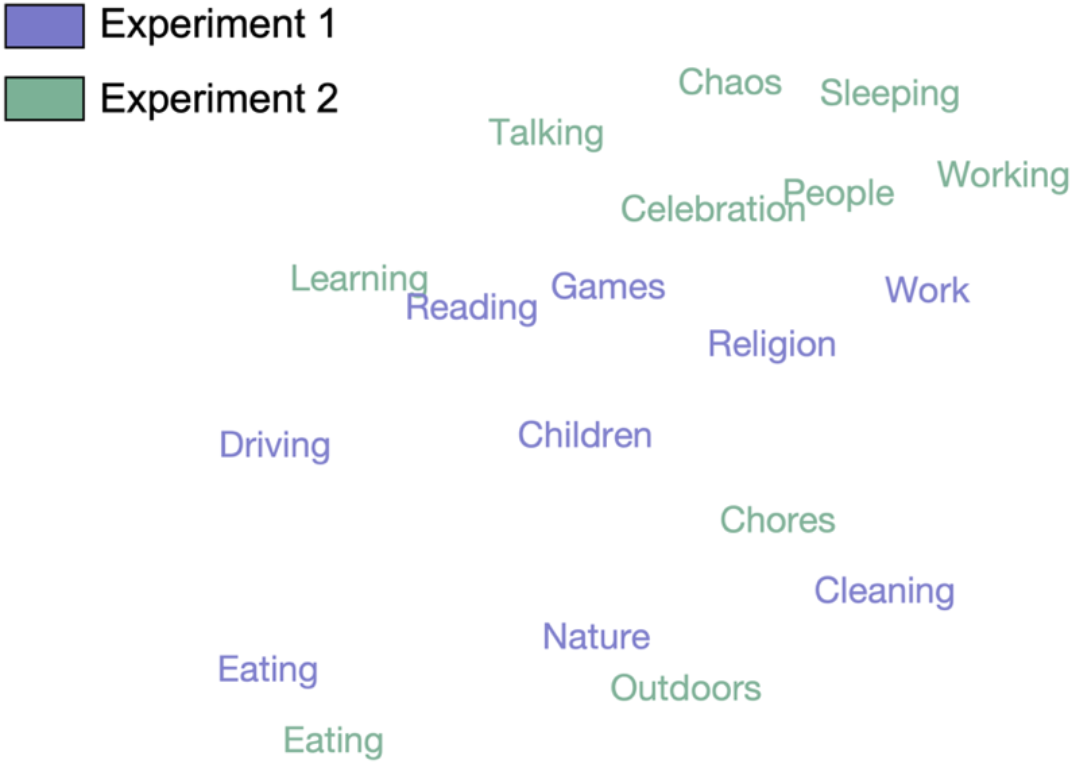
T-SNE plot displaying the distances between the averaged embeddings corresponding to each dimension from both experiments in a 2D space. *Eating, nature, cleaning, reading*, and *work* are the dimensions that most clearly replicate across experiments.

## Discussion

Here, we used sparse non-negative matrix factorization to recover a low-dimensional representation of intuitive action similarity judgments across two naturalistic video datasets. This resulted in robust and interpretable dimensions that generalized across experiments. Our results highlight the visual, semantic and social axes that organize intuitive visual action understanding.

### Non-negative matrix factorization as a viable approach to understanding similarity judgments

In the visual domain, it is reasonable to assume that features can be either absent or present to variable degrees, and that they can be additively combined to characterize a stimulus. Previous work has demonstrated that sparsity and positivity constraints enable the detection of interpretable dimensions underlying object similarity judgments (Hebart et al., 2020). Here, we showed that a different approach with the same constraints can recover robust, generalizable and interpretable dimensions of human actions. As opposed to those recovered for objects, the action dimensions were only moderately sparse, potentially due to the naturalistic nature of our stimuli, with many of them reflecting a combination of different dimensions. However, optimizing sparsity enabled us to strike the right balance between categorical and continuous descriptions of our data, thus capturing a rich underlying feature space (Hebart et al., 2020; Navarro & Lee, 2004; Zheng et al., 2019).

Our approach recovered a similar number of dimensions across the two experiments (ten and nine), despite their different stimulus set sizes (152 vs. 65 videos). While the dimensions all had an interpretable, semantic description, none mapped directly onto previously used visual, semantic, or social features, suggesting that a data-driven approach can uncover additional information beyond hypothesis-driven analyses. Furthermore, the dimensions generalized across important stimulus categories like action category and scene setting (Figure 2).

While a cross-validated PCA analysis uncovered a similar number of dimensions (eight), there was higher variance in the number and content of dimensions obtained after manipulating stimulus set composition (Supplementary Figures 4–5). Visual inspection of the dimensions also suggested that they may be less interpretable than those uncovered by sparse NMF. For example, two dimensions in Experiment 1 appeared to depict driving videos as the highest-weighted, yet these were interspersed with videos from different categories (e.g. cooking or socializing) that would make these dimensions difficult to label. The NMF *driving* dimension, on the other hand, showed the highest weights for the eight driving videos present in the dataset. Together, these results suggest that the positivity and sparsity constraints applied by NMF enable it to recover more robust and interpretable components from human behavioral data than PCA. These benefits are likely to extend to neural data, as suggested by the recent application of NMF to reveal novel category selectivity in human fMRI data (Khosla et al., 2022).

### Mapping internal action representations

We validated the resulting NMF dimensions in separate behavioral experiments. All dimensions were reproducible in an odd-one-out task (Figure 3A) and consistently labeled by participants, as quantified through semantic embeddings (Figure 3B). We visualized the most commonly assigned labels and assessed how they related to each other across the two experiments.

These analyses revealed several interpretable and reproducible dimensions, including those related to environment (*nature/outdoors*) and common everyday actions (*work*, *cleaning/chores*, *eating*, *reading/learning*). Importantly, these semantic distinctions emerged in the absence of a semantic task: participants were free to use their own criteria to define similarity, and the repetition of video pairs across multiple trials meant that different features could be used for judging similarity throughout the task. A previous data-driven analysis of semantic action similarity judgments found six clusters of actions related to locomotion, cleaning, food, leisure, and socializing (Tucciarelli et al., 2019). Here, we found that some semantic categories emerged in the absence of an explicit task, while other dimensions reflected visual or social-affective features. Indeed, several dimensions were given labels pertaining to people (*children/family, talking, people*), highlighting the social structure of the similarity data revealed by our previous hypothesis-driven work (Dima et al., 2022).

Importantly, the NMF procedure did not simply return the action categories used to curate the dataset, and in fact none of the dimensions provided a one-to-one correspondence with semantic action category (Figures 4–5). Instead, the dimension labels suggest that certain action categories were more salient than others (e.g. *work* or *eating*), while others tended to be grouped together based on other critical features. For example, activities that take place outdoors, like hiking and certain sports, were grouped together under a *nature/outdoors* dimension. In Experiment 1, this dimension included control videos depicting natural scenes, while in Experiment 2, this dimension emerged in the absence of such control videos, suggesting that the natural environment is a salient organizing feature in itself (Figure 4). In Experiment 2, videos depicting different actions were grouped together based on social or affective features like communication (talking face-to-face or on the phone) or negative affect (the *chaos* dimension, present, among others, in videos of people crying or fighting). This outcome is particularly important when dealing with naturalistic stimuli, where the actions depicted often belong to multiple categories (e.g. *talking+eating* or *fighting+driving*). In this case, a semantic model that does not take these complexities into account would fail to explain the behavioral data.

The dimension labels revealed differences as well as similarities between the two experiments. Notably, dimensions in Experiments 2 included more social-affective information (Figure 4), despite the fact that the two stimulus sets included the same action categories and were well-matched along social and affective dimensions (see Dima et al., 2022). However, the stimulus set in Experiment 2 was smaller, and stimulus sampling was conducted differently across the two experiments, resulting in more reliable similarity judgements in Experiment 2 (see Methods: Multiple arrangement). Despite these differences, the majority of dimensions correlated across experiments, suggesting that the NMF reconstructions form a shared semantic space, emerging in spite of stimulus set and sampling differences across experiments.

### From actions to event representations

How should action categories be defined? This is a challenging question, particularly given neuroimaging evidence that actions are processed in the brain at different levels of abstraction (Dima et al., 2022; Iordan et al., 2015; Spunt et al., 2016). Our results suggest that coarse semantic, visual, and social distinctions organize internal representations. The ACT-FAST taxonomy derived from data-driven text analysis proposed six broad distinctions (Thornton & Tamir, 2021a); however, our dimensions are more concrete and specific, likely reflecting our input of visually depicted everyday human actions. Two dimensions (*food* and *work*) emerged in both the text data and our two video datasets. This opens exciting avenues for research into visual and language-based action understanding and whether they share a conceptual taxonomy.

Relatedly, stimulus selection is the biggest factor in determining the structure of similarity judgments. Here, both stimulus sets represented 18 everyday action categories based on the American Time Use Survey, curated so as to minimize visual confounds. While the number of stimuli does not impact the dimensionality of the final NMF reconstruction, the number of action categories does (Supplementary Figure 3), and thus an accurate map of internal action representations will depend on comprehensive sampling of the relevant action space. Our results highlight a number of critical dimensions that organize how we judge the most common everyday actions; however, future research should expand this with datasets that sample actions in different ways, taking into account cultural and group differences in how we spend our time.

The low dimensionality of the NMF reconstruction may seem surprising. Actions bridge visual domains, including scenes, objects, bodies and faces, and thus vary along a wide range of features. Furthermore, our use of naturalistic videos adds a layer of complexity compared to previous work using still images. However, a low-dimensional internal representation is more likely to enable the efficient and flexible action recognition that guides human behavior.

Together, our results highlight the low-dimensional structure that supports human action representations, and open exciting avenues for future research. Our stimuli and the resulting dimensions bridge the boundary between actions and events, suggesting that our data-driven approach can be extended beyond specific visual domains to investigate how conceptual representations emerge in the mind and brain.

## Methods

### Stimuli

We analyzed two video datasets (Dima et al., 2022), each consisting of three-second naturalistic videos of everyday actions from the Moments in Time dataset (Monfort et al., 2019).

The videos were selected to represent the following 18 common action categories based on the American Time Use Survey (ATUS, 2019): childcare; driving; eating; fighting; gardening; grooming; hiking; housework; instructing; playing games; preparing food; reading; religious activities; sleeping; socializing; sports; telephoning; and working. The dataset used in Experiment 1 included 152 videos, with 8 videos per action category and 8 control videos depicting natural scenes or objects. The dataset used in Experiment 2 included 65 videos, with 3-4 videos per action category. For more details, see Dima et al., 2022.

### Participants

We analyzed data from two previously conducted multiple arrangement experiments (Dima et al., 2022). Experiment 1 involved 374 participants recruited via Amazon Mechanical Turk (300 after exclusions, located in the United States, gender and age not collected). 58 participants recruited through the Department of Psychological and Brain Sciences Research Portal at Johns Hopkins University took part in Experiment 2 (53 after exclusions, 31 female, 20 male, 1 non-binary, 1 not reported, mean age 19.38±1.09).

Two experiments were conducted to validate the dimensions resulting from Experiments 1 and 2. 54 participants validated the dimensions from Experiment 1 (51 after exclusions, 33 female, 13 male, 1 non-binary, 4 not reported, mean age 19.25±1.18) and a different set of 54 participants validated the dimensions from Experiment 2 (51 after exclusions, 37 female, 11 male, 3 not reported, mean age 20.12±1.78). All subjects were recruited through the Department of Psychological and Brain Sciences Research Portal at Johns Hopkins University.

All procedures for online data collection were approved by the Johns Hopkins University Institutional Review Board, and informed consent was obtained from all participants.

### Multiple arrangement

To measure the intuitive similarity between videos depicting everyday action events, we implemented a multiple arrangement task using the Meadows platform (www.meadows-research.com). Participants arranged the videos inside a circular arena according to their similarity. In order to capture intuitive, natural behavior, we did not define or constrain similarity. An adaptive algorithm ensured that different pairs of videos were presented in different trials, until a sufficient signal-to-noise ratio was achieved for each distance estimate. Behavioral representational dissimilarity matrices (RDM) were then constructed using inverse multi-dimensiomal scaling (Kriegeskorte & Mur, 2012). See Dima et al., 2022 for more details on the experimental procedure.

In Experiment 1, different subsets of 30 videos from the 152-video set were shown to different participants. The resulting behavioral RDM contained 11,476 video pairs with an average of 11.37±3.08 ratings per pair.

In Experiment 2, participants arranged all 65 videos. The resulting behavioral RDM contained 2,080 video pairs with 53 ratings per pair.

### Non-negative matrix factorization (NMF)

We used a data-driven approach, sparse NMF (Hoyer, 2002, 2004), to investigate the dimensions underlying action representations. This method has two important advantages over other forms of matrix decomposition, such as principal component analysis (PCA).

In aiming to represent each action video through a combination of underlying features, some of these may be assumed to be categorical. Such features would be present in some of the videos, but not in others, such that participants would arrange videos from the same category close together, and those outside the category farther apart. Sparse NMF applies sparsity constraints, allowing us to detect such categorical features that may group specific actions together.

However, the degree to which a feature is present may also distinguish certain actions from others, especially for features that capture non-categorical information. By enforcing positivity, NMF recovers continuous features with interpretable numerical values, reflecting the degree to which each feature is present in each stimulus. These two constraints thus allow both categorical and continuous structure to emerge, an approach well-suited to capture how real-world stimuli are represented in the mind (Navarro & Lee, 2004; Zheng et al., 2019).

Given a data matrix *V*, NMF outputs a basis vector matrix *W* and a coefficient matrix *H* with specified levels of sparsity and with *k* dimensions, such that *V ≈ WH*. Since NMF can output different results when initialized with random matrices, we used non-negative singular value decomposition for initialization (Boutsidis & Gallopoulos, 2008).

We first converted the behavioral RDM to a similarity matrix as used in symmetric applications of NMF (Kuang et al., 2012). As this matrix was symmetric, the output matrices were highly correlated (Pearson’s *r*>0.93), leading in practice to a similar solution to that given by symmetric NMF, where *W* = *H^T^*. We used nested cross-validation on ~90% of the data for parameter selection, with the rest of the data (9.52% in Exp 1 and 9.43% in Exp 2) held out to evaluate the final performance (Figure 1B).

In Experiment 1, cross-validation was implemented by leaving out one individual similarity rating for each pair of videos (orange in Figure 1B). Since different subjects had arranged different subsets of videos, this helped ensure sufficient data per pair of videos and cross-validation fold. The training similarity matrix was then created by averaging the remaining ratings for each pair. Any missing datapoints within a fold were imputed (no more than 0.2% of any given similarity matrix).

In Experiment 2, cross-validation was implemented across participants, and there were no missing datapoints. The final hold-out matrix included the data of five participants.

The two sparsity parameters for *W* and *H* were selected using two-fold cross-validation on two thirds of the training data. To speed up computation, we only tested combinations of sparsity parameters (*s*) ranging between 0 (no sparsity) and 0.8 (80% sparsity) in steps of 0.1. We selected the combination with maximal accuracy across the average of both folds, defined as the Kendall’s *τ_A_* correlation between the reconstructed *WH* matrix and the test matrix. This analysis was repeated for different numbers of dimensions (*k*) up to 150 in Experiment 1 and 65 in Experiment 2 (just below the maximum number of videos in each experiment). The best combination of sparsity parameters was applied to the remaining one third of the training data.

Five iterations of the above cross-validation for parameter selection were performed, and the best number of dimensions was selected using the average performance curve on the held-out training set. To avoid overfitting, we identified the elbow point in this performance curve, defined as the point maximally distant from a line linking the two ends of the curve.

The NMF procedure was then reinitialized with the output of the first cross-validation fold and rerun on the whole training set (90% of the data) with the selected combination of parameters. The held-out 10% of the data was used to evaluate performance.

#### Control analyses relating NMF dimensions to stimulus categories

We performed a post-hoc control analysis to assess the robustness of NMF dimensions to perturbations in the stimulus set. The NMF procedure was repeated after leaving out key stimulus categories that correlated with identified NMF dimensions (outdoors, indoors, childcare, driving, and fighting). To ensure these stimulus categories did not drive results, the dimensions obtained from each control analysis were correlated to the original dimensions. The correlations were then tested against chance using one-tailed randomization testing with 1000 iterations of component matrix shuffling.

To evaluate whether NMF dimensions captured any obvious stimulus features (e.g. scene setting, action category or sociality), we assessed the correlation between each NMF dimension and 12 visual, action-related, and social features (Supplementary Figure 2; Dima et al., 2022).

#### Control PCA analysis

To asssess whether NMF provides an advantage over the more commonly used PCA, we conducted a similar cross-validated analysis using PCA, and assessed the resulting reconstruction accuracy and robustness to stimulus set perturbations in both experiments. The cross-validation procedure was exactly the same, except that no search for sparsity parameters was conducted. Instead, only the number of dimensions (*k*) was selected using two-fold cross-validation on the training data (~90% of the data).

### Dimension validation

We used two tasks in two separate online experiments (corresponding to Experiments 1 and 2) to assess the interpretability of NMF dimensions in separate participant cohorts. We presented the eight highest weighted and eight lowest weighted videos along each dimension obtained from NMF as stimuli to the subjects. The experiment was implemented in JavaScript.

First, participants were asked to select the odd video out of a group consisting of seven highly weighted videos and one low-weighted video (odd-one-out) for a given dimension. This was done 20 times for each dimension with random resampling (from the top and bottom eight) of the videos shown. Participants were excluded if they did not achieve above-chance performance (over 12.5%) on catch trials involving a natural scene video as the odd-one-out among videos containing people. Dimensions were considered reproducible if participants achieved above-chance accuracy in selecting the odd-one-out (sign permutation testing, 5000 iterations, omnibus-corrected for multiple comparisons).

After completing this task, participants were asked to provide up to three labels (words or short phrases) for each dimension based on a visual inspection of the eight highest and eight lowest weighted videos.

#### Semantic analyses

We visually inspected the labels provided by participants to correct spelling errors and identify cases where pairs of antonyms were used to label a dimension (e.g. *nature vs home);* in these cases, we only kept the first label. Next, we visualized the labels by creating word clouds of the most common labels using the MATLAB *wordcloud* function.

To quantify participant agreement on labels, we used FastText (Bojanowski et al., 2016), a 300-dimensional word embedding pretrained on 1 million English words. Embeddings were generated for each of the words and phrases provided by participants. Euclidean distances were then calculated across all labels within each dimension. Labels were considered related if the distance between them was in the 10^th^ percentile across dimensions and experiments (below a threshold of *d* = 1.2). To generate a chance level for participant agreement, we calculated the proportion of related labels across different dimensions.

Finally, we assessed whether the NMF dimension labels replicated across the two experiments. To generate a dissimilarity matrix, embeddings were averaged across labels within each dimension before calculating Euclidean distances between dimensions. This allowed us to visualize which dimensions were most semantically related across experiments.

## Supporting Information

**Supplementary Figure 1.**
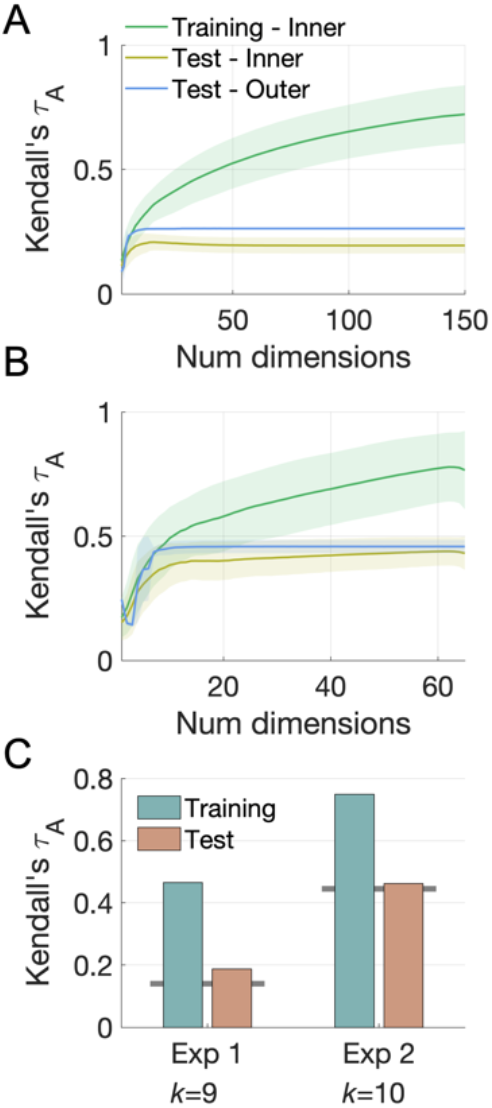
NMF performance. **A.** Training set NMF reconstruction performance in Experiment 1 evaluated on the three sections of the training set (see Figure 6). Error bars are ± 1SD. Performance on the outer set in the nested cross-validation procedure plateaus with nine dimensions. **B.** As in A, for Experiment 2. **C.** Final performance of the reconstructed matrix on the whole training set (~90% of the data) and the held-out set (~10% of the data), plotted against the true hold-out correlation (gray horizontal bars).

**Supplementary Figure 2.**
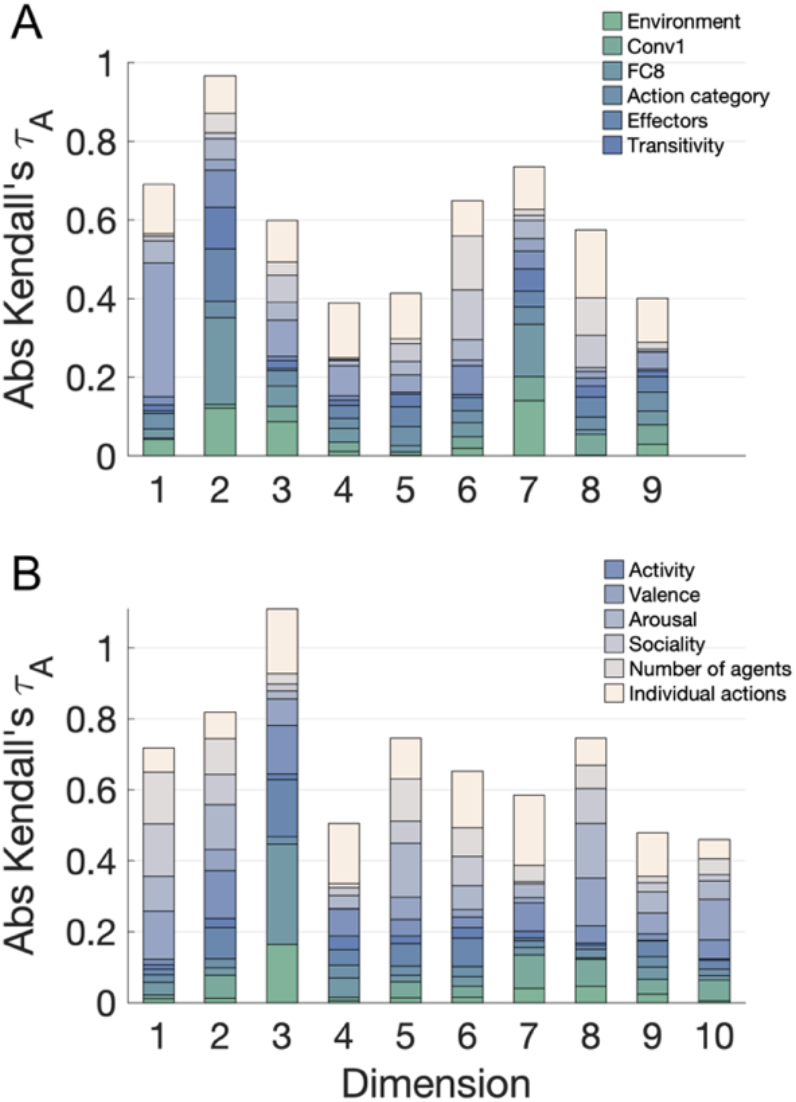
Absolute correlations between visual, social, and action features and each NMF dimension for **A.** Experiment 1 and **B.** Experiment 2, shown in a stacked plot. Dimension are sorted in descending order of their summed weights.

**Supplementary Figure 3.**
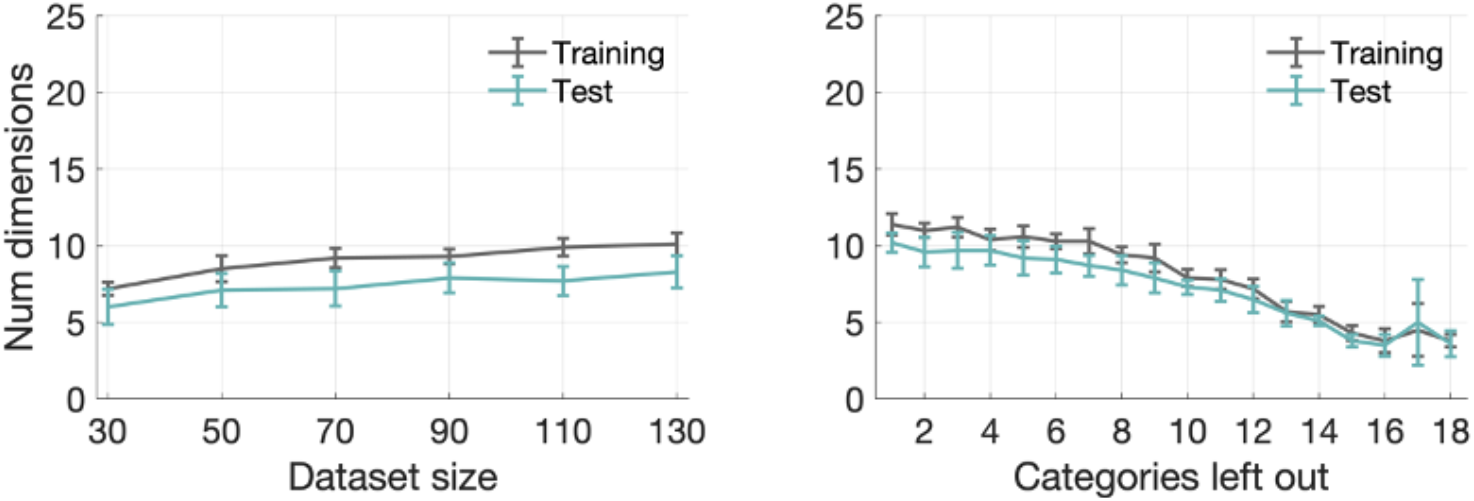
Dimensionality as a function of dataset size. **Left**: number of dimensions obtained by running the NMF procedure using random subsets of the stimuli from Experiment 1 (10 iterations). **Right**: number of dimensions obtained by running NMF after leaving out a number of randomly selected action categories (10 iterations). The number of action categories has a larger effect on dimensionality than the dataset size.

**Supplementary Figure 4.**
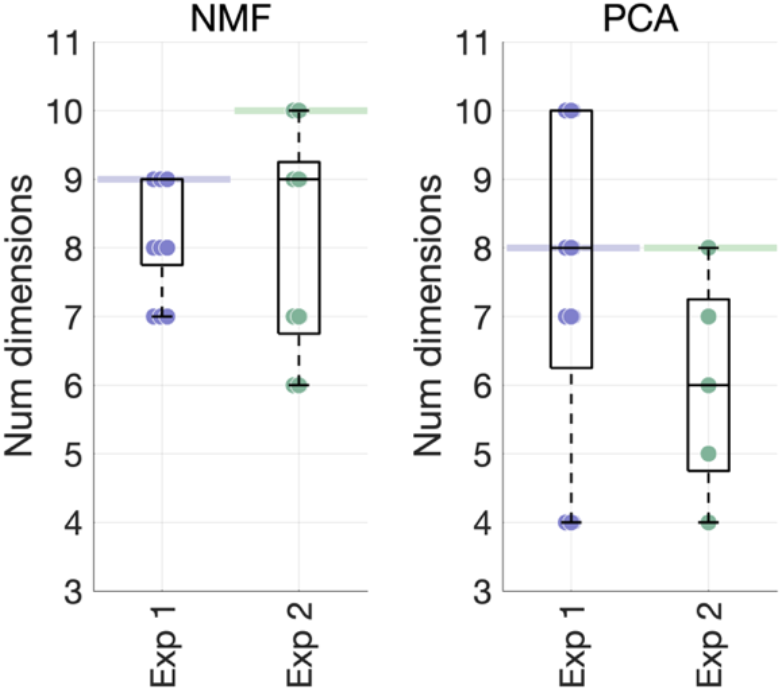
NMF and PCA dimension robustness. The PCA procedure was repeated five times, after removing key stimulus categories from the behavioral RDM from Experiment 1. Each dot shows the number of dimensions resulting from each iteration, with horizontal lines showing the number of dimensions recovered from the full datasets.

**Supplementary Figure 5.**
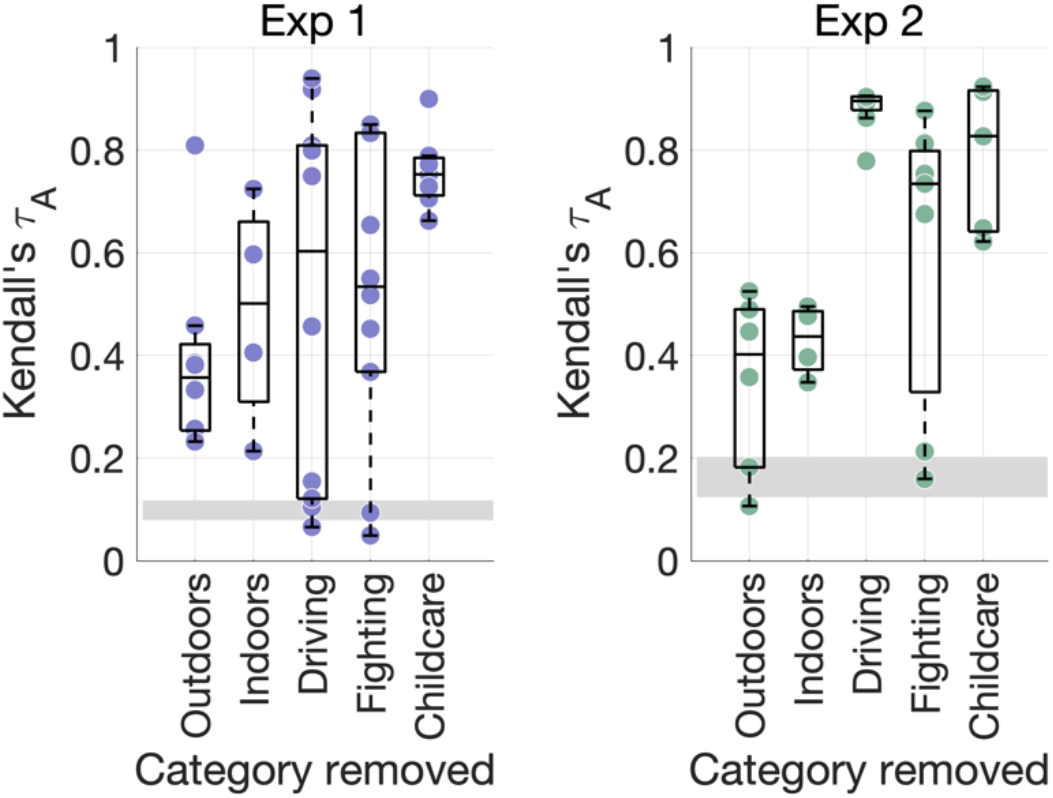
PCA dimension robustness. The PCA procedure was repeated five times, after removing key stimulus categories from the behavioral RDM from Experiment 1. Each dot shows the maximal correlation between each dimension obtained in the control analysis and any of the original dimensions with the same stimuli removed (repeats allowed). The grey rectangles depict the chance level. Although on average correlations are higher than those obtained with NMF, their variance is overall almost twice as high, suggesting that stimulus set perturbations have a stronger impact on some of the PCA dimensions.

**Supplementary Figure 6.**
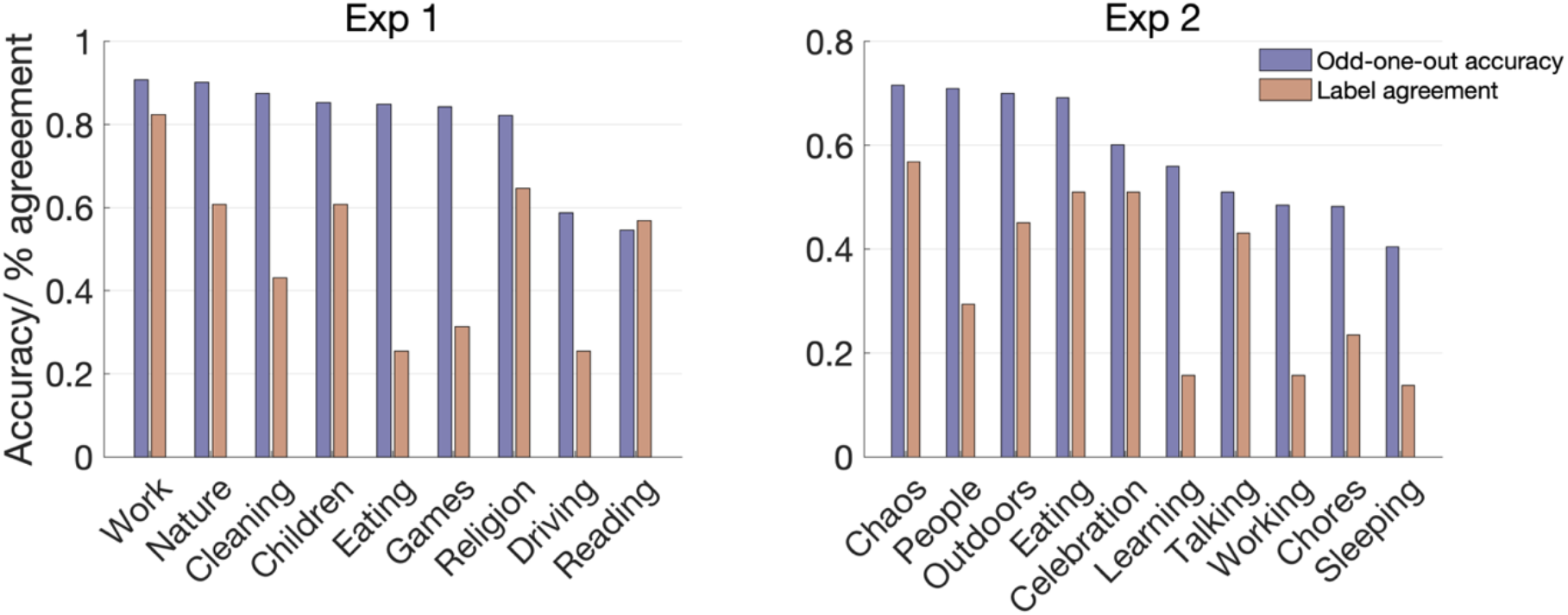
Validation variability. Dimensions are named according to their most common labels, and ranked according to the accuracy obtained for each of them in the odd-one-out task. Participant agreeement on the most common label is also shown for each dimension.

**Supplementary Figure 7.**
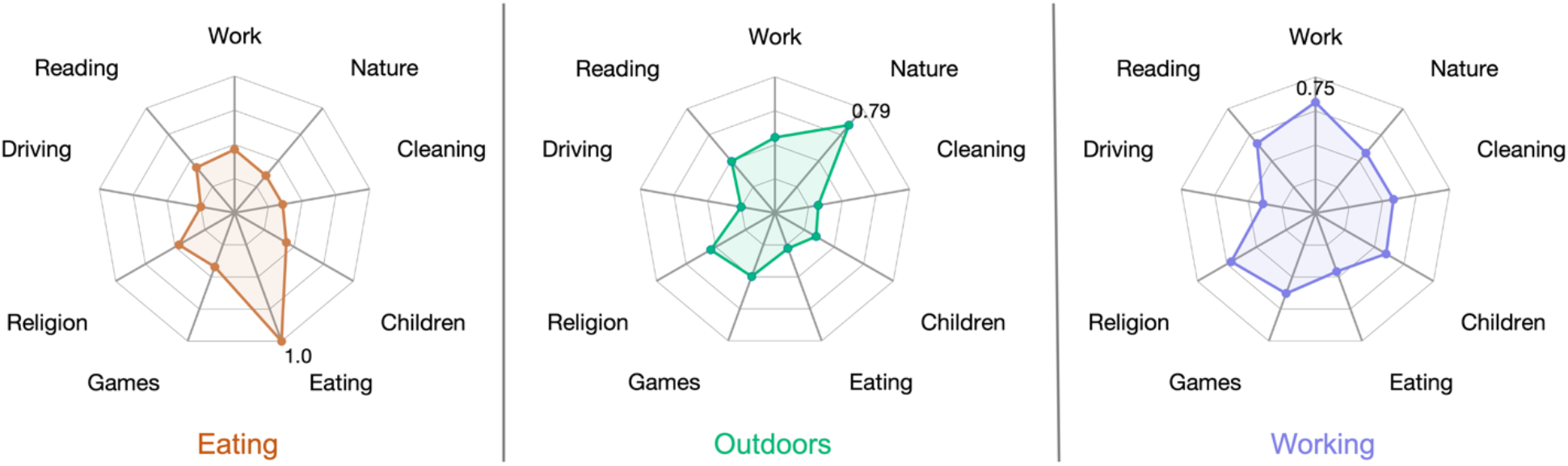
Examples of how three dimensions from Experiment 2 map onto the dimensions from Experiment 1, as measured via semantic embeddings of the labels given by participants. The similarity values shown are relative (i.e. normalized to the 0-1 range).

